## Supplementary materials for "Modified MRI anonymization (de-facing) for improved MEG coregistration"

### Supplementary Material

#### *CBU group: ratings of familiarity*

The CBU group was familiar with some of the people whose images were used. Individual familiarity ratings provided by the CBU group were positively related to the identification accuracy at the single-trial level. This was confirmed in a linear mixed effects logistic regression that modelled condition, order, and familiarity as fixed effects, and subject and faces as random effects. The analysis revealed a significant linear effect of familiarity ($F(1,1474) = 3.73$, $p < .05$) in addition to a significant condition by order interaction ($F(6,1474) = 2.48$, $p < .05$).

#### *Face identification effect*

Some faces were easier to identify than others, most likely due to some idiosyncratic features in their face and head shape. An Analysis of Variance (ANOVA) confirmed a significant main effect of Face ($F(9,810$, epsilon=0.87) = 22.2, $p < .001$) when collapsing across defacing conditions, but this effect was also observed in analysis that solely focussed on the defaced condition ($F(9,810$, epsilon=0.81) = 6.48, $p < .001$) see Figure S1 and S2.

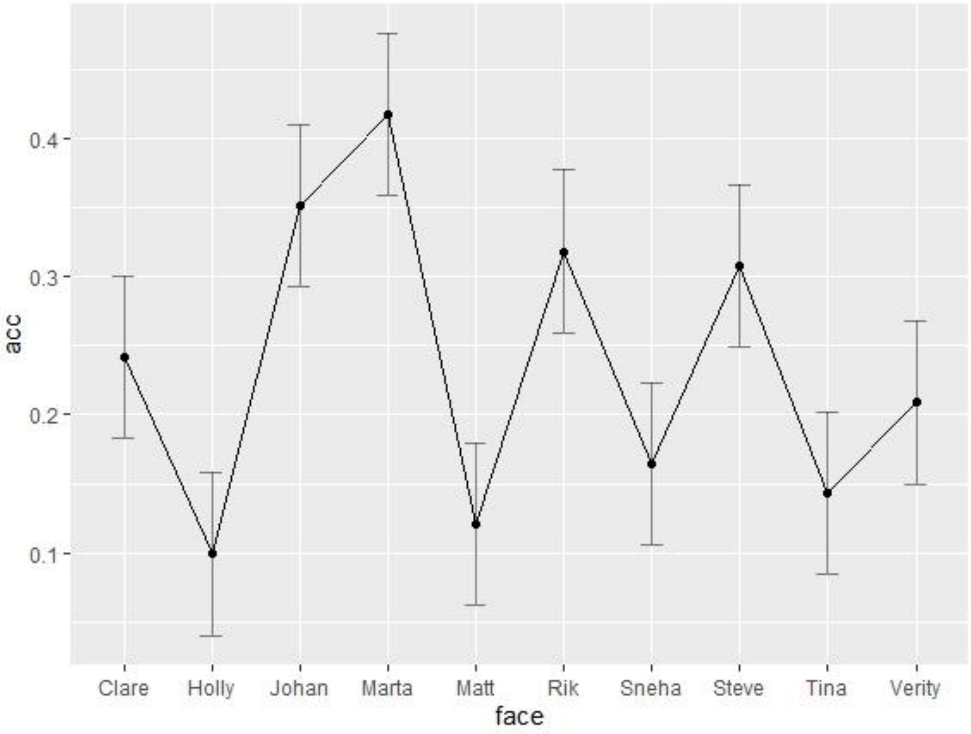


Fig. S1A: Accuracy in face identification was systematically better for some faces compared to others, even when considering the Defaced condition only, which is shown here collapsed across different blocks.

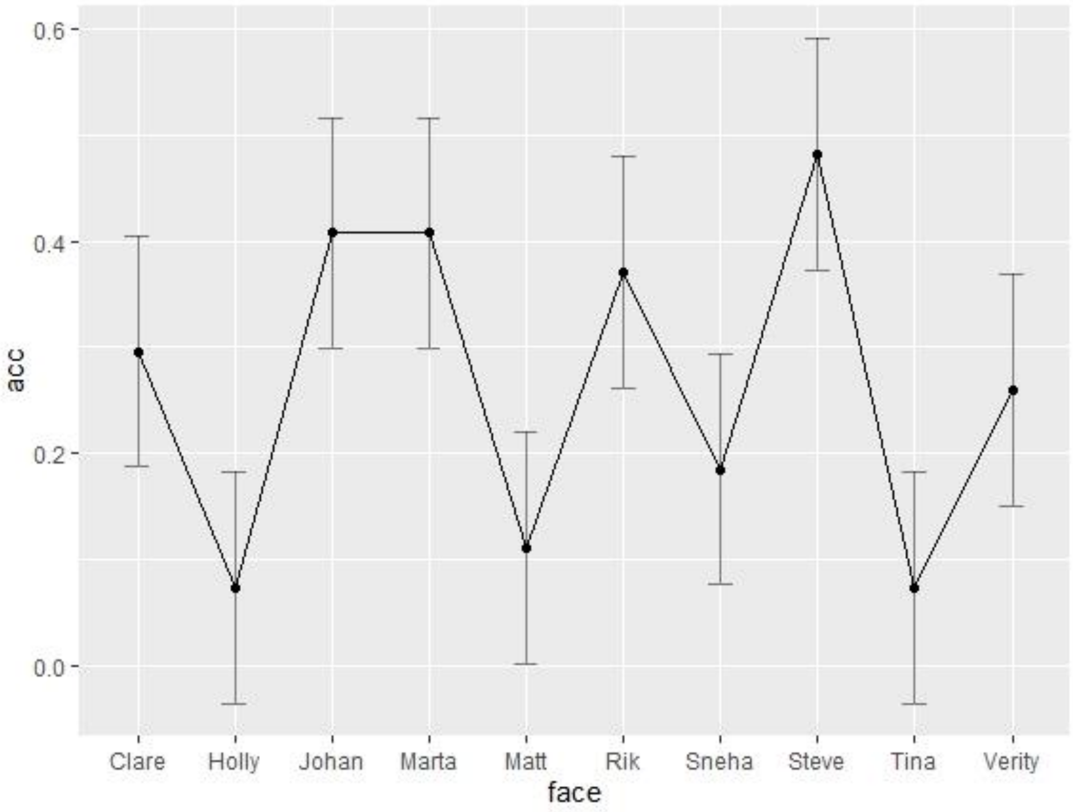


Fig. S1B: Accuracy in face identification was systematically better for some faces compared to others, even when considering the Defaced condition only, now shown for the first block only.
